# Nifedipine Modulates Renal Lipogenesis via the AMPK-SREBP Transcriptional Pathways

**DOI:** 10.1101/487264

**Authors:** Yen-Chung Lin, Mai-Szu Wu, Chang-Rong Chen, Chang-Yu Chen, Chang-Jui Chen, Kuan-Chou Chen, Chiung-Chi Peng

## Abstract

Lipid accumulation in renal cells has been implicated in the pathogenesis of obesity-related kidney disease, and lipotoxicity occurring in the kidney can be a surrogate marker for renal failure or renal fibrosis. Nifedipine-induced renal lipotoxicity has never been cited, although a few studies have shown that nifedipine inhibits lipogenesis via activation of the LKB1-AMPK pathway. Therefore, we utilized NRK52E cell models to examine this further. We pre-treated cells with varying concentrations of nifedipine (7.5, 15, or 30 μM) for 24 or 48 h prior to examining the activity of lipogenesis enzymes and lipotoxicity.

Nifedipine was found to activate acetyl CoA synthetase, acetyl CoA carboxylase, long chain fatty acyl CoA elongase, ATP-citrate lyase, and HMG CoA reductase, suggesting elevated production of cholesterol, triacylglycerides, and phospholipids. Nifedipine exposure induced a vast accumulation of cytosolic free fatty acids (FFA) and stimulated the production of reactive oxygen species, upregulated CD36 and KIM-1 (kidney injury molecule-1) expression, inhibited p-AMPK activity, and triggered the transcription of *SREBP-1/2* and *lipin-1*, underscoring the potential of nifedipine to induce lipotoxicity with renal damage.

In the present study, we elucidated the mechanism of action of nifedipine that leads to renal lipotoxicity via the AMPK-SREBP-1/2 pathway. To our knowledge, this is the first report demonstrating nifedipine-induced lipid accumulation in the kidney.

## Introduction

Ectopic lipid deposition in non-adipose tissues such as the kidney may elicit organ damage with subsequent lipotoxicity, further affecting the function of the involved organs [1]. Overproduction of free fatty acids (FFA) may enhance the translocation of FFA into mitochondria, triggering the production of reactive oxygen species (ROS) in the early phase of accumulation. This results in increased mitochondrial proton conductance and uncoupling of oxidative phosphorylation, eventually provoking mitochondrial permeabilization [2], which underlies dysfunction in fatty acid oxidation (FAO). Inhibition of FAO in renal proximal tubule epithelial cells causes depletion of ATP, cell death, de-differentiation, and intracellular lipid deposition, all of which contribute to the phenotype of renal fibrosis [3].

When kidney cells are injured, individual cell types are differently affected. The endothelial cells undergo apoptosis, inducing inflammation and mesangial cell proliferation and eventual glomerulopathy. Furthermore, podocyte apoptosis and ER stress may be evoked, resulting in proteinuria and glomerulopathy. Lastly, the tubular cells also undergo apoptosis with increased autophagy vesicles, leading to interstitial inflammation and fibrosis [4].

CD36 [cluster of differentiation 36, also known as platelet glycoprotein 4, fatty acid translocase (FAT) or scavenger receptor B2] is a scavenger receptor that functions in high-affinity tissue uptake of long-chain fatty acids (LCFAs) and, under conditions of excessive fat supply, contributes to lipid accumulation and metabolic dysfunction [5]. Renal CD36 is mainly expressed in the proximal tubule cells, podocytes, and mesangial cells, and is markedly upregulated in the setting of chronic kidney disease (CKD) [6]. Recently, growth hormone releasing peptides (GHRP) have been identified as potent inducers of PPARγ and its downstream actions through the activation of CD36, demonstrating an alternative method of regulating essential aspects of hepatic cholesterol biosynthesis and fat mitochondrial biogenesis [7].

Hepatic lipogenesis has been reported to be mediated predominantly by the AMPK/SREBP-1 pathway in rat hepatocytes and human hepatoma cell lines [8]. A diet incorporating oleic acid (OA) decreased the phosphorylation of AMPK and increased the maturation of SREBP-1 and the expression of SREBP-responsive genes [8]. Mice and mouse hepatocytes were resistant to OA-induced lipogenesis because of little if any response by AMPK and downstream effectors [8].

An increasing number of studies have linked the activation of sterol regulatory element-binding proteins (SREBPs), the dysfunction of peroxisome proliferator-activated receptors (PPARs), and inhibited β-oxidation of fatty acids with diabetic nephropathy [9]. High glucose may initiate lipid deposition on renal tubular cells by upregulating CD36 and PPARs [10]. SREBP-1c, regulated by the insulin and AMPK signaling pathways, was found to play a role in nonalcoholic fatty liver disease [11]. In addition, lipin plays an important role in lipid metabolic homeostasis [12]. Lipin-1 has been shown to act as a key mediator of the effects of mTORC1 on SREBP-dependent fatty acid and cholesterol biosynthesis [13].

It has been well documented that many thiazide-type diuretic antihypertensives exhibit a hyperlipidemic effect, while such adverse effects are seldom associated with the calcium channel blockers (CCBs) [14,15]. Research suggests that CCBs may impair renal autoregulation [16], and clinically, they may be less effective than other antihypertensives [17]. Moreover, the safety of CCBs in patients with proteinuria and renal insufficiency is widely questioned [18]. We hypothesize that nifedipine may promote renal damage due to its lipogenic effect, which is associated with the AMPK-SREBP-lipin pathway. In this present study, using the normal rat proximal tubular epithelial cell line NRK52E as the model, we investigated whether the lipogenesis of nifedipine is associated with the AMPK-SREB-lipin pathway.

## Materials and Methods

### Cell culture

The normal rat kidney epithelial-derived cell line NRK52E (CRL-1571) was obtained from the Bioresource Collection and Research Center, Food Industry Research Development Institute, Hsinchu, Taiwan. NRK52E cells were cultured in 5% bovine calf serum-supplemented Dulbecco’s modified eagle medium from Gibco (Carlsbad, CA, USA) at 37ºC in a humidified atmosphere with 5% CO_2_. Upon reaching 80% confluence, the cells were trypsinized with 0.25% trypsin–0.02% ethylenediaminetetraacetic acid (EDTA) for 5 min at 37ºC and then repassaged.

### Cell survival assay

NRK52E cells (2×10^4^ cells/well) were incubated in a 24-well plate for 16 h, and then treated with the nifedipine (Sigma, St. Louis, MO, USA) at different concentrations (0, 7.5, 15, or 30 μM) or with 150 μM oleic acid from Sigma (St. Louis, MO, USA) as a positive control for 48 h. The nifedipine dose was selected to correspond to internal levels in humans, and the positive control oleic acid dose was selected according to a preliminary 3-(4,5-dimethylthiazol-2-yl)-2-5-diphenyltetrazolium bromide (MTT) test. The cells were washed with phosphate-buffered saline (PBS) and incubated with 0.5 mg/mL MTT solution for 4 h at 37ºC prior to removing the culture medium. Dimethyl sulfoxide was then added and mixed for 5 min at 26ºC. Cell viability was determined by measuring the absorbance at 560 nm. Cell viability for each experimental group was calculated as a percentage of that of the control group.

### Oil Red O staining

NRK52E cells (2×10^4^ cells/well) were incubated in a 24-well-plate for 16 h, and then treated with nifedipine (0, 7.5, 15, or 30 μM) or 150 μM oleic acid for 24 h and 48 h, respectively. The cells were then fixed on the plate by 10% formalin for 3 h, the plate was washed with 60% isopropanol to dry, and then 300 μL of 3 mg/mL Oil Red O from Sigma (St. Louis, MO, USA) was added to the cells and left for 1 h to air dry. The plate was washed with flowing water and Oil Red O dye, and photographs were acquired. Finally, 200 μL of 100% isopropanol was added to extract residual Oil Red O, and the absorption was monitored at 515 nm.

### Lipid assay

NRK52E cells (2 × 10^4^) were incubated in 24-well plates, and then treated with nifedipine (0, 7.5, 15, or 30 μM) or oleic acid (150 μM) for 48 h. The lipid assay was conducted using the Total Cholesterol and Cholesteryl Ester Colorimetric/Fluorometric Assay Kit from Biovision (Milpitas, California, USA, #K603-100).

### Lipid peroxidation

The measurement of thiobarbituric acid reactive substances (TBARS) is the most widely employed assay to determine lipid peroxidation by measuring the production of malondialdehyde, which is a naturally occurring byproduct of lipid peroxidation. In this study, we evaluated the extent of lipid peroxidation using a standardized TBARS assay kit (Cayman Chemical, Ann Arbor, Michigan USA), according to the manufacturer’s instructions. Cell suspensions were centrifuged at 161 x g in a fixed angle rotor for 5 min and washed twice with PBS. The supernatants were discarded, and the cell pellets were resuspended in 1 mL of PBS and sonicated on ice. The malondialdehyde thiobarbituric acid adduct formed by the reaction was measured colorimetrically at 530 ~ 540 nm.

### Measurement of ROS production

Dihydroethidium (DHE) was detected by flow cytometry. NRK52E cells (2 × 10^4^) were incubated in 24-well plates and then treated with nifedipine (30 μM) for 4 h or with H_2_O_2_ (500 μM) for 30 min as the positive control. The cells were dyed with the Muse Oxidative Stress Kit (MCH100111; Millipore, Billerica, MA, USA) and subjected to flow cytometry to determine the degree of DHE production. ROS flow cytometry was repeated at least three times.

### Western blotting procedure

A standard western blotting protocol was used as described previously [19,20]. The primary antibodies used in this study included the Fatty Acid and Lipid Metabolism Antibody Sampler Kit (#8335), anti-TNFR1 (#13377), anti-AMPK (#5832), and anti-p-AMPK (#2535) from Cell Signaling Technology (Danvers, MA, USA); anti-SREBP-1 (sc-13551) and anti-SREBP-2 (sc-13552) from Santa Cruz Biotechnology (Santa Cruz, CA, USA); anti-CD36 (NB400-144ss), anti-β actin (NB600-501), and anti-α tubulin (NB100-690, Novus Biologicals, Littleton, CO, USA); anti-PPARα (GTX01098), anti-HDAC1 (GTX100513), and anti-histone H3 (GTX122148) from GeneTex, Irvine, CA, USA); and anti-KIM 1 (ab190696, Abcam, Cambridge, MA, USA). The secondary antibody used was either anti-mouse IgG or anti-rabbit IgG (Jackson ImmunoResearch, West Grove, PA, USA), which was dissolved in 5% skim milk in TBST for 1 h, followed by incubation for 1–2 min in enhanced chemiluminescence mixture (JT96-K004M, T-Pro Biotechology, Zhonghe, New Taipei City, Taiwan) for visualization. The western blot was repeated at least three times.

### Statistical analysis

Analysis of variance and post-hoc tests using SPSS 14.0 (SPSS Inc., Chicago, IL, USA) were used to statistically evaluate the data, and results are presented as the mean ± standard deviation. Two-tailed *p*-values ≤ 0.05 (marked as **) were considered statistically significant. In addition, *p*-values ≤ 0.01 (marked as ***) are denoted.

## Results and Discussion

### Effect of nifedipine on cell viability

Nifedipine dose- and time-dependently suppressed the viability of NRK52E cells (Fig 1a). After incubation for 24 h and 48 h, nifedipine at 7.5 μM did not show any effect on the viability of NRK52E cells. At 15 μM and 30 μM, nifedipine suppressed cell viability to 15.6% and 39.0% relative to the control and 40.3% of that induced by oleic acid (*p* < 0.01) (Fig 1a) during the first 24 h. The cell viability was further suppressed to 24.4% and 47.2% relative to the control after 48 h, compared with 42.3% caused by oleic acid (*p* < 0.01) (Fig 1 a).

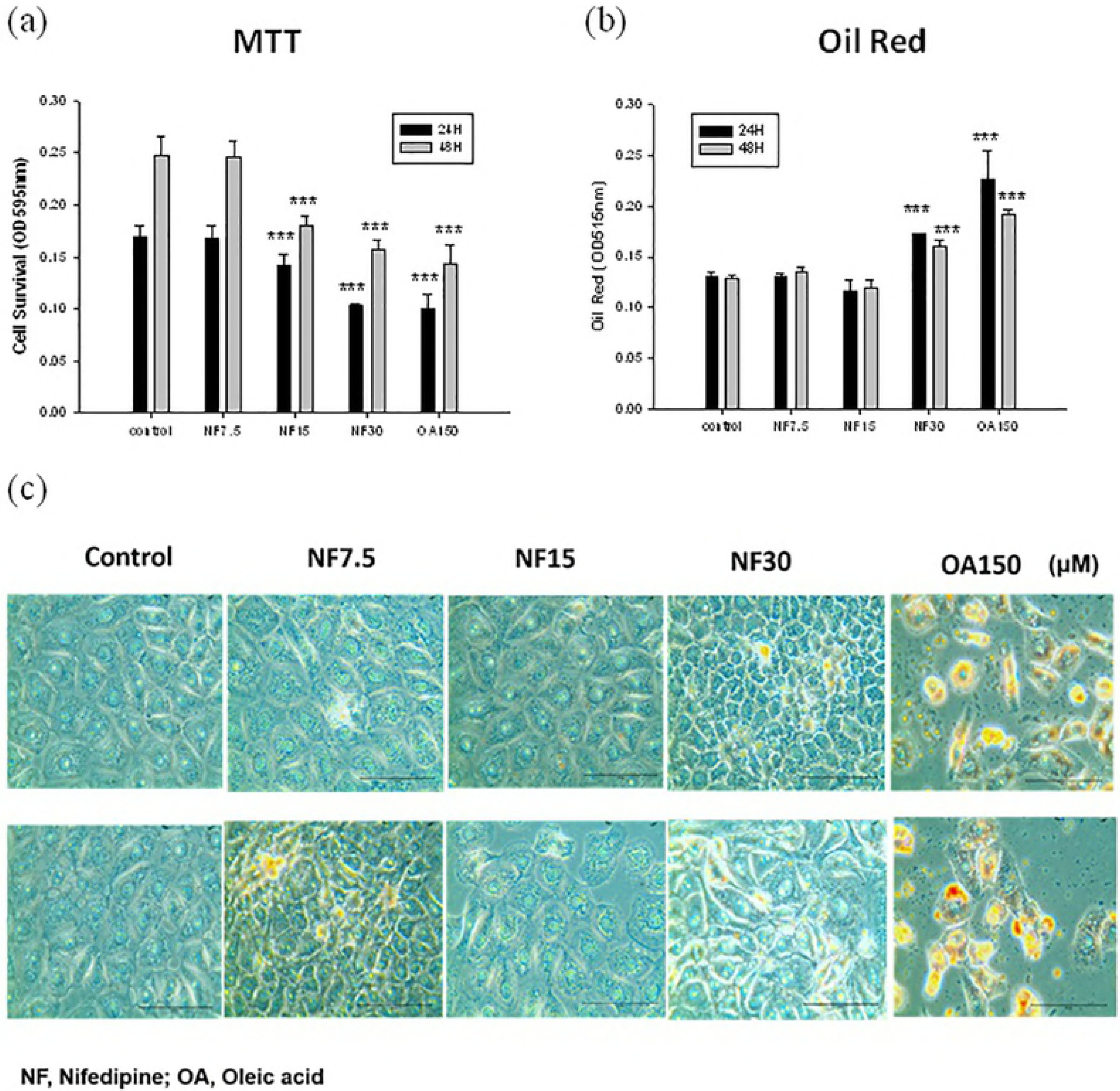

### Effect of nifedipine on intracellular lipid accumulation

Oil Red staining revealed that the intracellular lipid content significantly increased with increasing doses of nifedipine (7.5 µM vs. 30 µM, *p* < 0.01) (Fig 1b, Fig 1c). Moreover, it is worth mentioning that the cell size shrank in a dose-dependent manner starting from the original spindle-fat shape (control), with slight swelling at 7.5 μM, a larger triangular shape at 15 μM, and finally a granular polygonal shape at 30 µM (upper panel of Fig 1c). The change became more apparent after incubation for 48 h (lower panel of Fig 1c), suggesting the possibility that the NRK52E cells could have been affected by nifedipine therapy both morphologically and physiologically and that the deviation of the cytoplasmic lipid accumulation at 15 μM could be caused by such cellular physiological changes.

### Nifedipine stimulated production of reactive oxygen species (ROS)

Nifedipine at 30 μM stimulated the production of ROS at a 3.3-fold rate compared to the control group in ROS (+)/ROS(-) after treatment with nifedipine for 16 h, whereas a 2.57-fold increase was observed after treatment with 500 μM H_2_O_2_ for 30 minutes (*p* < 0.01) (Fig 2a).

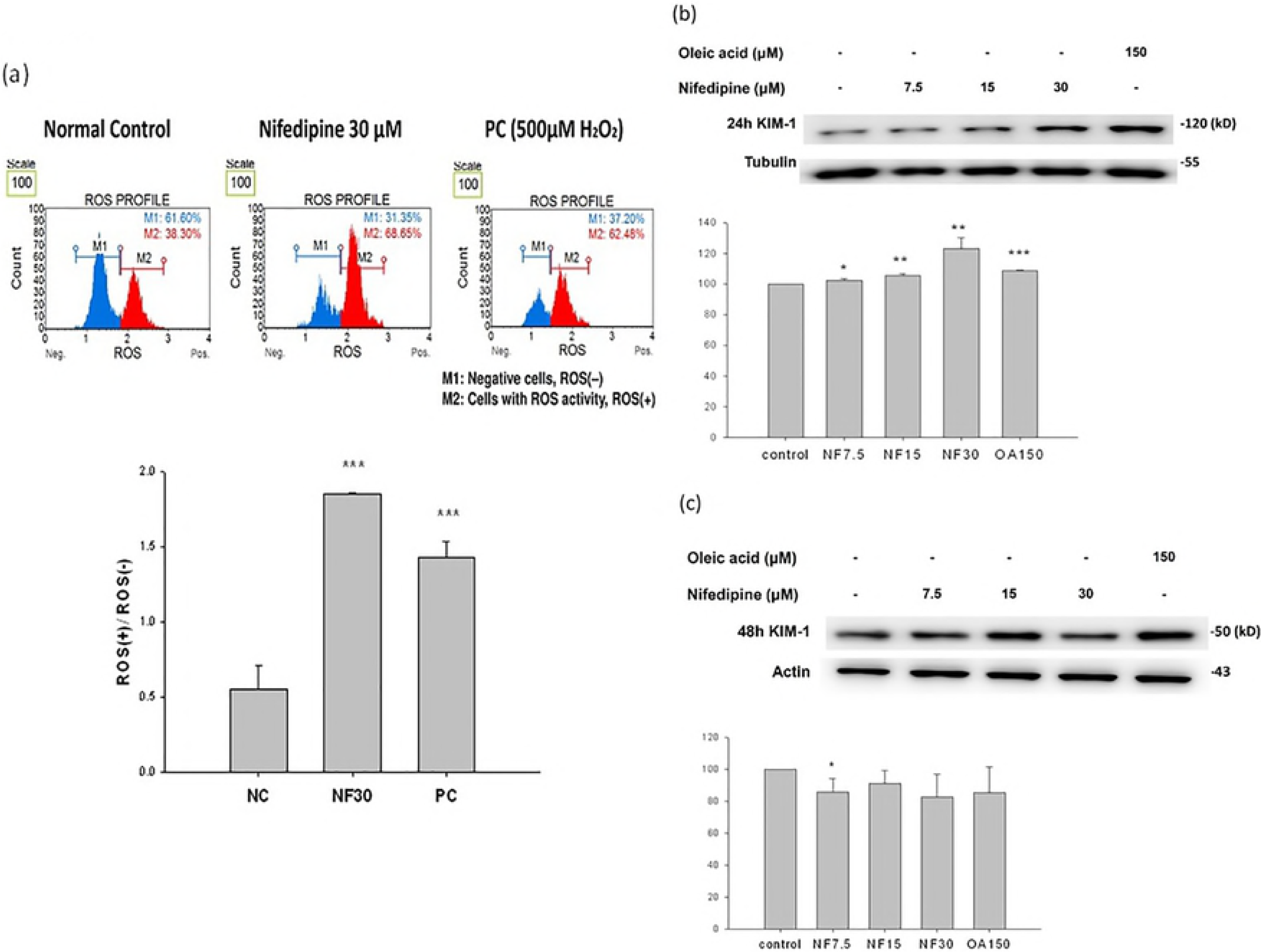

### Nifedipine promoted the expression of proteins related to kidney injury

The expression of KIM (kidney injury molecule-1; also known as T cell immunoglobulin mucin domains–1, TIM-1) was simultaneously upregulated in dose dependent fashion to 101%, 102%, and 122% (*p* < 0.01) compared to control, respectively, following nifedipine at 7.5, 15, and 30 μM for 24 h, compared with 103% induced by oleic acid (Fig 2b) and then decreased to 86.0%, 91.0%, and 80% at 48 h (Fig 2c) related to control. KIM-1 is not detectable in the normal human and rodent kidney, but its expression increases more than that of any other protein in the injured kidney, and it is localized predominantly to the apical membrane of the surviving proximal epithelial cells [21].

In humans, blood KIM-1 levels are significantly elevated in the setting of acute kidney injury (AKI) and chronic kidney disease (CKD) and predicted progression of renal disease in a type 1 diabetic cohort [22], suggesting a possible link between nifedipine therapy and kidney damage. There are many reasons why KIM-1 may be released into the circulation after kidney proximal tubule injury. In kidneys with injury, the tubular cell polarity is lost, such that KIM-1 may be released directly into the interstitium. Further, increased transepithelial permeability after tubular injury leads to a backleak of tubular contents into the circulation [23]. Humphreys et al. demonstrated that chronic KIM-1 expression led to inflammation and tubule interstitial fibrosis, characterized by elevated monocyte chemotactic protein-1 (MCP-1) levels and increased MCP-1-dependent macrophage chemotaxis [24].

In experimental AKI, the intensity of KIM-1 expression increased in proportion to the severity of injury and was consistently present in segment S3 (the collecting tubule, mostly cortical, mainly representing the proximal straight tubule), but only transiently in other segments (i.e. segments S1 and S2, proximal convoluted tubule). Vimentin was absent in the proximal tubules of healthy cats but expressed in injured S3. These findings indicate that S3 is the proximal tubular segment most susceptible to ischemic injury and that KIM-1 is a sensitive tissue indicator of AKI in cats [25]. The upregulation of KIM-1 at 24 h with rapid decline at 48 h may implicate the status of AKI.

### Nifedipine stimulated cholesterol *de novo* biosynthesis

Nifedipine stimulated *de novo* cholesterol biosynthesis in a non-dose- and non-time-dependent manner. The cholesterol level reached a range of 111.3 μg/mL (nifedipine 7.5 μM) to 114.9 μg/mL (nifedipine 30 μM), compared with 93-5-97.4 μg/mL for the control) in 24 h or 48 h, respectively (*p* < 0.01) (Fig 3a).

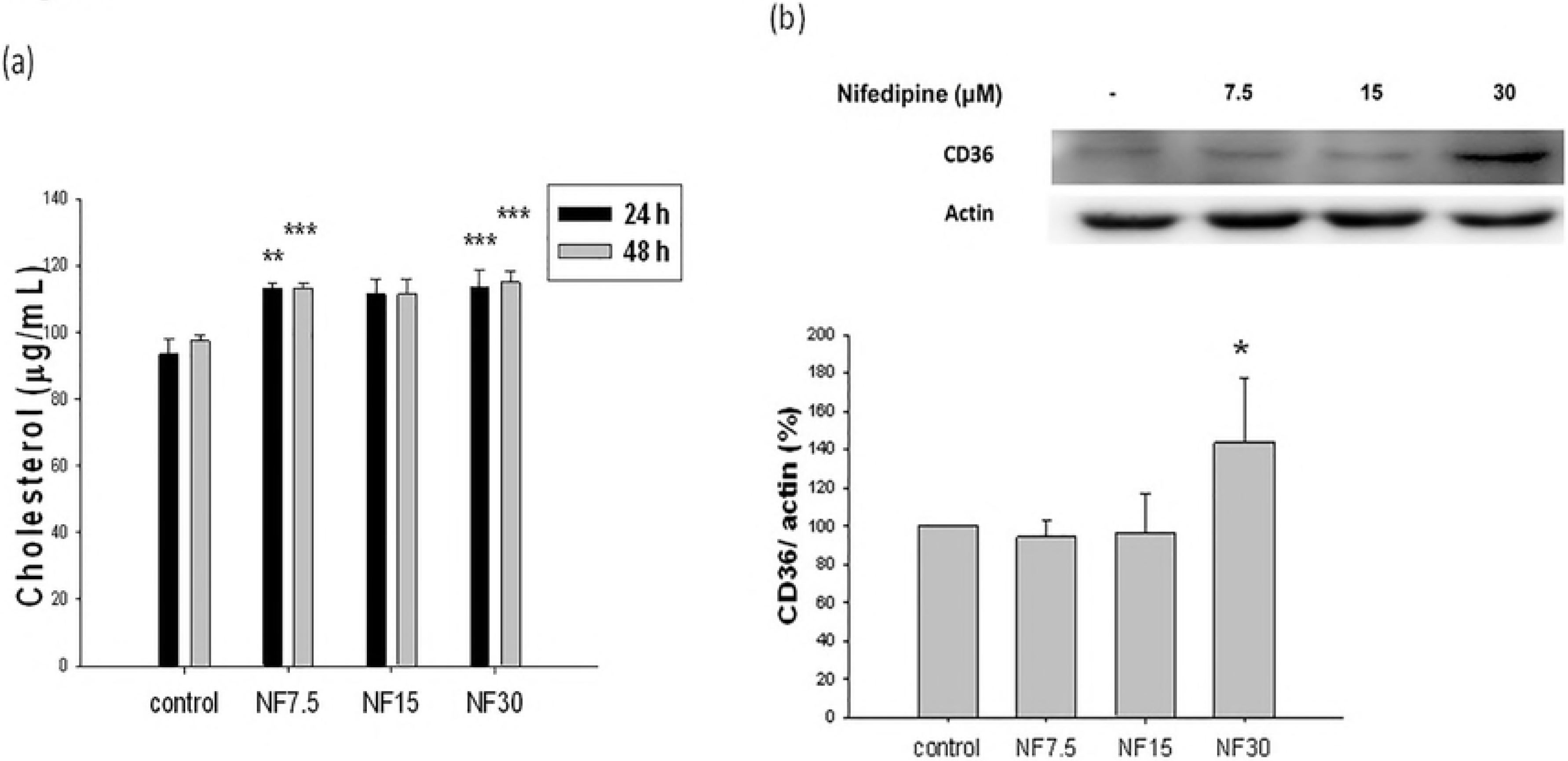

### Nifedipine upregulated the expression of CD36 proteins related to lipid translocation

CD36 was upregulated by nifedipine at 30 μM to 143.6 % (*p* < 0.1), but unaffected at the lower doses 7.5 μM and 15 μM (Fig 3b). In addition to acting as a fatty acid translocase, CD36 also functions as a novel mediator influencing binding and uptake of albumin in the proximal tubule, and CD36 is upregulated in proteinuric renal diseases. Proximal tubular CD36 expression was markedly increased in proteinuric individuals [26]. Hyperlipoproteinemia and its subsequent oxidation are usually associated with glomerular capillary dysfunction and severe glomerulosclerosis in dyslipidemic patients due to lipid deposits in glomeruli [27, 28]. Diet-induced hypercholesterolemia led to renal endothelial dysfunction associated with vascular and microvascular remodeling, inflammation, and kidney fibrosis [29]. Thus, the upregulation of CD36 induced by nifedipine (30 μM) in the rat kidney NRK52E cell line supports the hypothesis that nifedipine at regular therapeutic dosages could provoke renal cholesterol accumulation (Fig 3a, Fig 3b).

Palmitic acid and high-fat diets cause lipotoxicity *in vivo* and *in vitro* and adversely switch the energy source from the CD36 pathway to the GLUT4 pathway [12]. Recently, a review by Maréchal et al. has emphasized that growth hormone releasing peptides (GHRP) are a potent inducer of PPAR through activation of CD36 [7], thereby regulating essential aspects of lipid and energy metabolism [30].

To understand whether the cholesterol accumulating effect of nifedipine was due to stimulation of the *de novo* biosynthetic pathway or merely because of translocation, we further explored the effect of nifedipine on the *de novo* biosynthetic pathway of cholesterol.

### Nifedipine stimulated the production of acetyl CoA synthetase (ACS), acetyl CoA carboxylase (ACC), and long chain fatty acyl elongase (ACSL1), but not fatty acid synthase (FAS) and p-ACC

Nifedipine stimulated the production of the enzyme acetyl CoA synthase (ACS) (*p* < 0.05 at 7.5 μM; *p* < 0.01 at 15 μM and 30 μM compared to controls) (Fig 4a), acetyl CoA carboxylase (ACC) to 128.5% and 144.4% of the control values at 15 μM and 30 μM doses (*p* < 0.05) (Fig 4b), and long chain fatty acyl elongase (ACSL1) (*p* < 0.01 at all concentrations) (Fig 4c), but simultaneously inhibited fatty acid synthase (FAS) (Fig 4d), and p-ACC (*p* < 0.01) because of inhibitory phosphorylation of ACC (Fig 4b). Two pathways of butyrate synthesis by fatty acid synthetase have been identified [31]. Prior research has shown that increasing the malonyl-CoA concentration and the ratio of malonyl-CoA to acetyl-CoA concentrations increased the proportion of long-chain (C_14:0_–C_18:0_) and decreased the proportion of short-chain (C_4:0_ and C_6:0_) fatty acids formed [31], with long-chain fatty acids synthesized mainly as free acids and short-chain fatty acids occurring predominantly in the esterified form. Medium-chain fatty acids (C_8:0_–C_12:0_) were formed in amounts approximately equimolar to the fatty acid synthetase protein [31]. In our case, the concentration of malonyl CoA might be sufficiently high, but the ratio of malonyl CoA/acetyl CoA could be very low due to the high concentration of acetyl CoA produced from citrate by action of ATP citrate lyase, leading to suppressed fatty acid synthase.

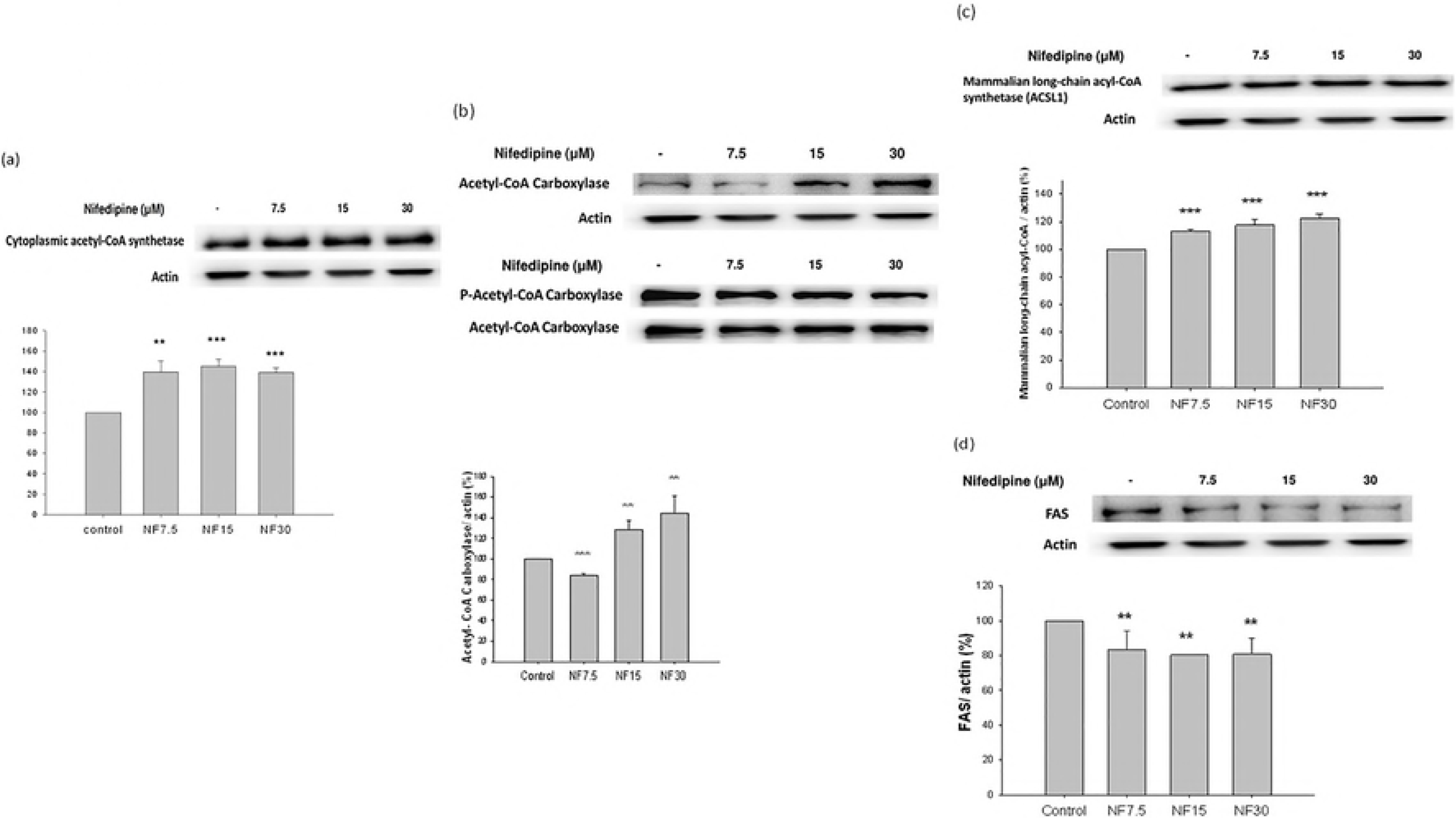

Other studies have demonstrated that nifedipine and amlodipine prevented non-esterified fatty acids (NEFA)-induced endothelial dysfunction, leukocyte activation, and enhancement of oxidative stress without affecting blood pressure [32]. As most NEFA are present in form of FFA, we speculate that nifedipine could have inhibited the production of FAS.

### Nifedipine stimulated the production of ATP citrate lyase (ACL) and HMG-CoA reductase

The amount of ATP citrate lyase produced was stimulated by nifedipine in a dose dependent manner, with 7.5 μM and 15 μM doses increasing production to 150% and 169%, and then leveling off at 30 μM at 170%, compared with the control (*p* < 0.01) (Fig 5a). In parallel, the amount of downstream HMG-CoA reductase was also upregulated by nifedipine administration in a dose-dependent fashion to 107% and 120% at 15 μM and 30 μM doses as compared with the control (*p* < 0.01) (Fig 5b). This then led to increased *de novo* cholesterol biosynthesis (Fig 3a-b).

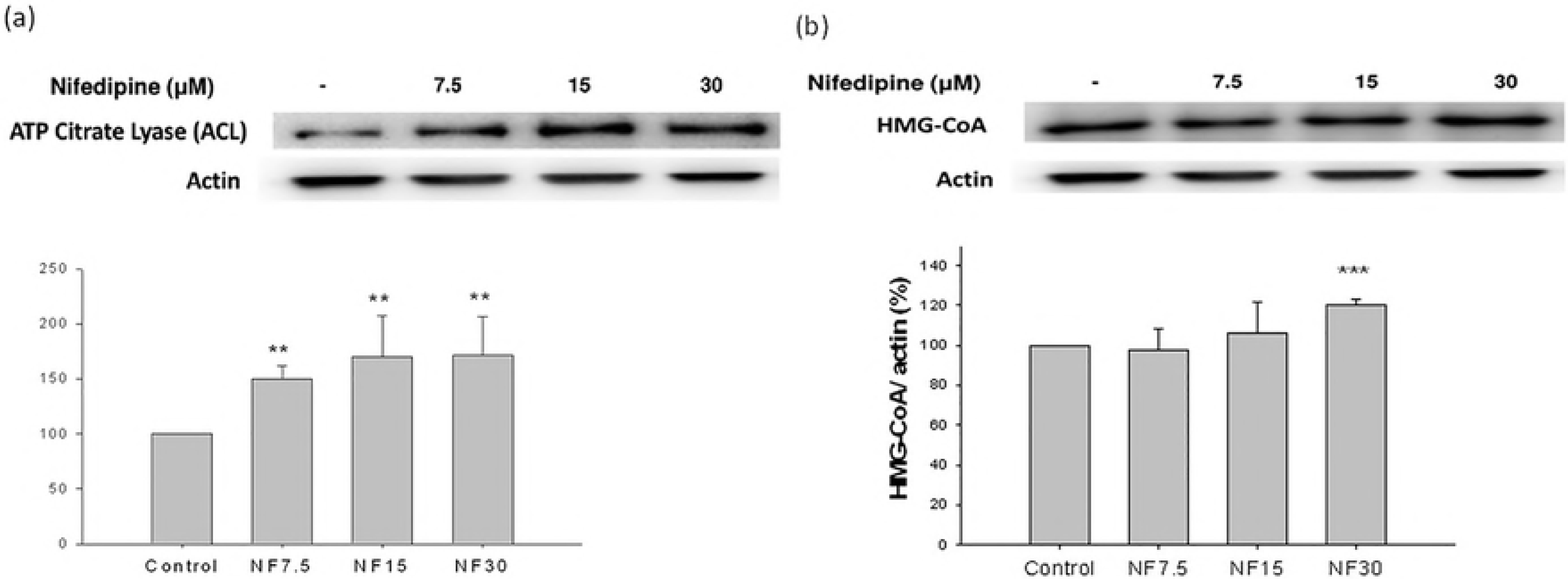

### Nifedipine downregulated the expression of phosphorylated AMPK

Although the level of 5′ AMP-activated protein kinase (AMPK)/actin was slightly yet significantly upregulated in a dose-dependent fashion following treatment with nifedipine to 107.1%, 112.6%, and 116.4% of control levels, more significantly, the ratio of p-AMPK to AMPK was downregulated in a dose-dependent manner to 82.1%, 58.7%, and 49.9% of control levels, respectively, at 7.5 μM, 15 μM, and 30 μM doses of nifedipine (Fig 6a). It is known that downregulation of AMPK upregulates mTOR and subsequently stimulates lipogenesis and adipogenesis, resulting in enhanced lipid storage [33]. Maintenance of the energy balance depends on the efficiency of tightly regulated mechanisms of energy intake and expenditure [34].

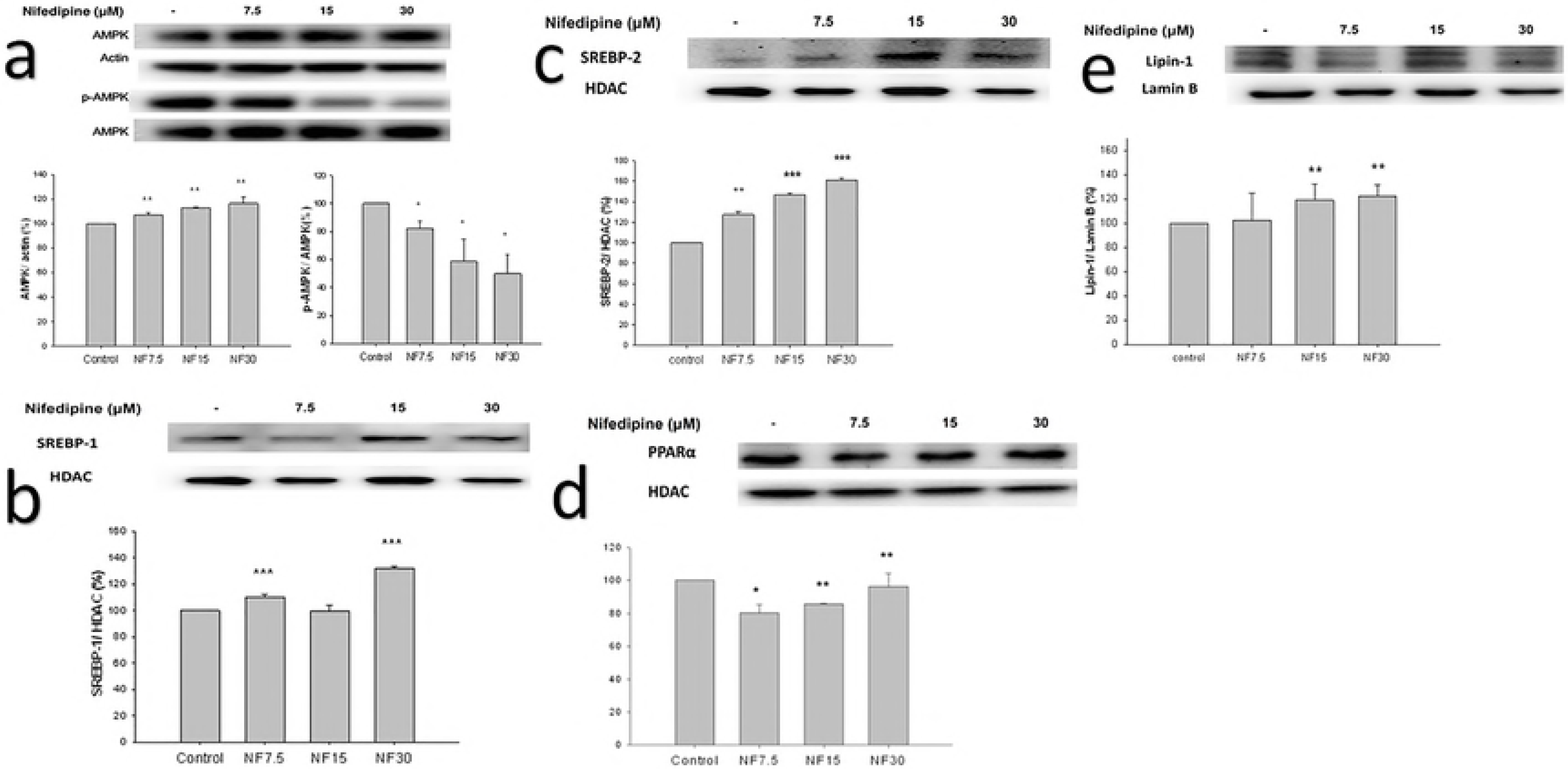

Establishing the correlation of a high-fat diet with diabetes and the expression of AMPK has been challenging. High-fat diet fed mice demonstrated a marked increase in markers of fibrosis and inflammation, with significantly suppressed AMPK in kidneys [35]. AMPK regulates NFκB activation and acts as a potent regulator of NADPH oxidase as well as the TGF-β system [36]. The activity of AMPK correlates with both obesity-related and diabetes-related renal diseases [36]. Activated AMPK reduced renal hypertrophy. Supposedly, the lipid accumulation in kidneys induced by nifedipine can inhibit the activation of AMPK (Fig 6a), and consequently may provoke early renal damage as caused by a high fat diet [35].

On the other hand, activated AMPK will phosphorylate and inactivate ACC, resulting in a decrease in the intracellular level of malonyl-CoA, thereby relieving inhibition of CPT-1 activity and accelerating lipolysis [37]. Conversely, downregulated p-AMPK/AMPK (Fig 6a) is expected to lead to an increased level of malonyl CoA, promoting the biosynthesis of triglycerides and phospholipids [38].

### Nifedipine upregulated the expression of SREBP-1 and SREBP-2

Sterol-regulatory element-binding proteins (SREBPs) are key transcription factors regulating the expression of a diversity of enzymes required for endogenous cholesterol, fatty acid, triglyceride, and phospholipid biosynthesis. [39] SREBP activity is tightly regulated to maintain lipid homeostasis and is modulated by extracellular stimuli such as growth factors.

The levels of SREBP-1/HDAC and SREBP-2/HDAC were upregulated following nifedipine treatment. Relative to controls, the SREBP-1/HDAC level was upregulated to 109.9% and 132.0% at 7.5 and 30 μM doses of nifedipine, respectively (*p* < 0.01) (Fig 6b); those of SREBP-2 were elevated to 127.5%, 146.7% and 161.0% of control levels by nifedipine at 7.5, 15, and 30 μM doses, respectively (Fig 6c). SREBP-1a, SREBP-1c and SREBP-2 act differently in lipid synthesis. SREBP-1c is involved in FA synthesis and insulin-induced glucose metabolism (particularly in lipogenesis), whereas SREBP-2 is relatively specific to cholesterol synthesis [39]. The SREBP-1a isoform is associated with both pathways [39]. The upregulation of both SREBP-1 and SREBP-2 (Fig 6b-c) suggests that both de novo lipogenesis and cholesterol synthesis. Consistent with this, the de novo biosynthesis of long chain fatty acid, triglycerides (Fig. 4), and cholesterol (Fig 3a, Fig 5) were all stimulated as evidenced by the upregulated ACC and ACSL1 (Fig 4), as well as ATP citrate lyase and HMG-CoA reductase (Fig 5).

In SREBP processing, the inactive precursors of SREBP transcription factors are synthesized bound to the endoplasmic reticulum (ER) membranes, and their function is mainly controlled by the cellular sterol content. When sterol levels decrease, the precursor is cleaved to activate cholesterogenic genes and maintain cholesterol homeostasis [39]. This sterol-sensitive process appears to be a major point of regulation for the SREBP-1a and SREBP-2 isoforms [39]. The unique activation properties of each SREBP isoform facilitate the coordinated regulation of lipid metabolism; however, further studies are needed to understand the detailed molecular pathways that specifically regulate each SREBP isoform [39].

### Nifedipine upregulated the expression of PPAR-α

Compared with the control group, the level of peroxisome proliferator-activated receptor-α (PPARα)/lamin B was suppressed at a low dose of nifedipine and then dose-dependently restored from 80.2%, 85.6%, to 96.4% upon administration of nifedipine at doses of 7.5, 15, and 30 μM, respectively (Fig 6d).

PPARs are ligand-activated transcription factors of the nuclear hormone receptor superfamily comprising the following three subtypes: PPARα, PPARγ, and PPARβ/δ.[40] PPARs are involved in various DNA-independent and DNA-dependent molecular and enzymatic pathways in adipose tissue, liver, and skeletal muscle. The PPAR family of nuclear receptors plays a major regulatory role in energy homeostasis and metabolic function [40]. Activation of PPAR-α reduces triglyceride levels and is involved in regulation of energy homeostasis. These pathways are affected in disease conditions and can cause metabolic energy imbalance [40].

### Nifedipine upregulated the expression of lipin-1

The level of lipin-1 was highly upregulated after nifedipine treatment to 102.5%, 118.9%, and 112.2% of control levels by 7.5, 15, and 30 μM of nifedipine, respectively (Fig 6e).

Lipins are cytosolic phosphatidate phosphatases (PAPs) involved in the glycerolipid biosynthesis pathway [41]. Proteins in the lipin family play a key role in lipid synthesis due to their PAP activity, and they also act as transcriptional coactivators to regulate the expression of genes involved in lipid metabolism [42].

Lipins in the nucleus act as transcriptional co-activators with peroxisome proliferator-activated receptor γ co-activator-1α (PPAR-γ C-1α; or PGC-1α), PPAR-α, and other factors such as histone acetyltransferase (HAT) to stimulate the expression of genes involved in fatty acid oxidation [43,44]. The lipin proteins usually reside in the cytosol. Translocation of lipin to the endoplasmic reticulum (ER) membrane occurs in response to elevated fatty acid levels. There, it catalyzes the conversion of phosphatidic acid (PA) to diacylglycerol (DAG), building the key substrates for the synthesis of triacylglycerol (TAG), phosphatidylethanolamine (PE), and phosphatidylcholine (PC) [42]. The increased expression of lipin-1 together with the fatty acid-induced translocation of lipin-1 and lipin-2 to the ER facilitates increased TAG synthesis in starvation, diabetes, and stress conditions [43].

The upregulation or downregulation of lipin-1 depends on the disease model [41]. Inhibition of hepatic mTORC1 significantly impairs SREBP function and makes mice resistant, in a lipin-1 - dependent fashion, to the hepatic steatosis and hypercholesterolemia induced by a high-fat diet [41]. In this study, we showed that nifedipine upregulated lipin-1 (Fig 6e), ACS (Fig 4a), ACC (Fig 4b), ACSL1 (Fig 4c), and FAS (Fig 4d), and dose-dependently enhanced the response to Oil Red (Fig 1b-c), indicating a strongly upregulated pathway of TAG and increased phospholipid biosynthesis. In addition, nifedipine stimulated *de novo* cholesterol biosynthesis (Fig 3a), as evidenced by highly upregulated ACL (Fig 5a), and HMG-CoA reductase (Fig. 5b). We speculate that lipin-1 could have been translocated from the cytosol to ER as previously mentioned [42]. More importantly, we have demonstrated that nifedipine upregulated SREBP-1 and SREBP-2 (Fig 6b-c) and, at lower doses, nifedipine suppressed PPAR-α (Fig 6d). Specifically and selectively, SREBP-1a activates fatty acid and cholesterol synthesis; SREBP-1c, fatty acid synthesis; and SREBP-2, cholesterol synthesis and uptake [45]. These data, taken together, strongly suggest that the lipogenic effect of nifedipine may be associated with the acute renal damage caused by renal lipotoxicity, as evidenced by the upregulation of KIM-1 (Fig 2b-c). Lipotoxicity is considered to be a surrogate marker for renal failure or renal fibrosis. A graphic summary of the results is shown in Fig 7.

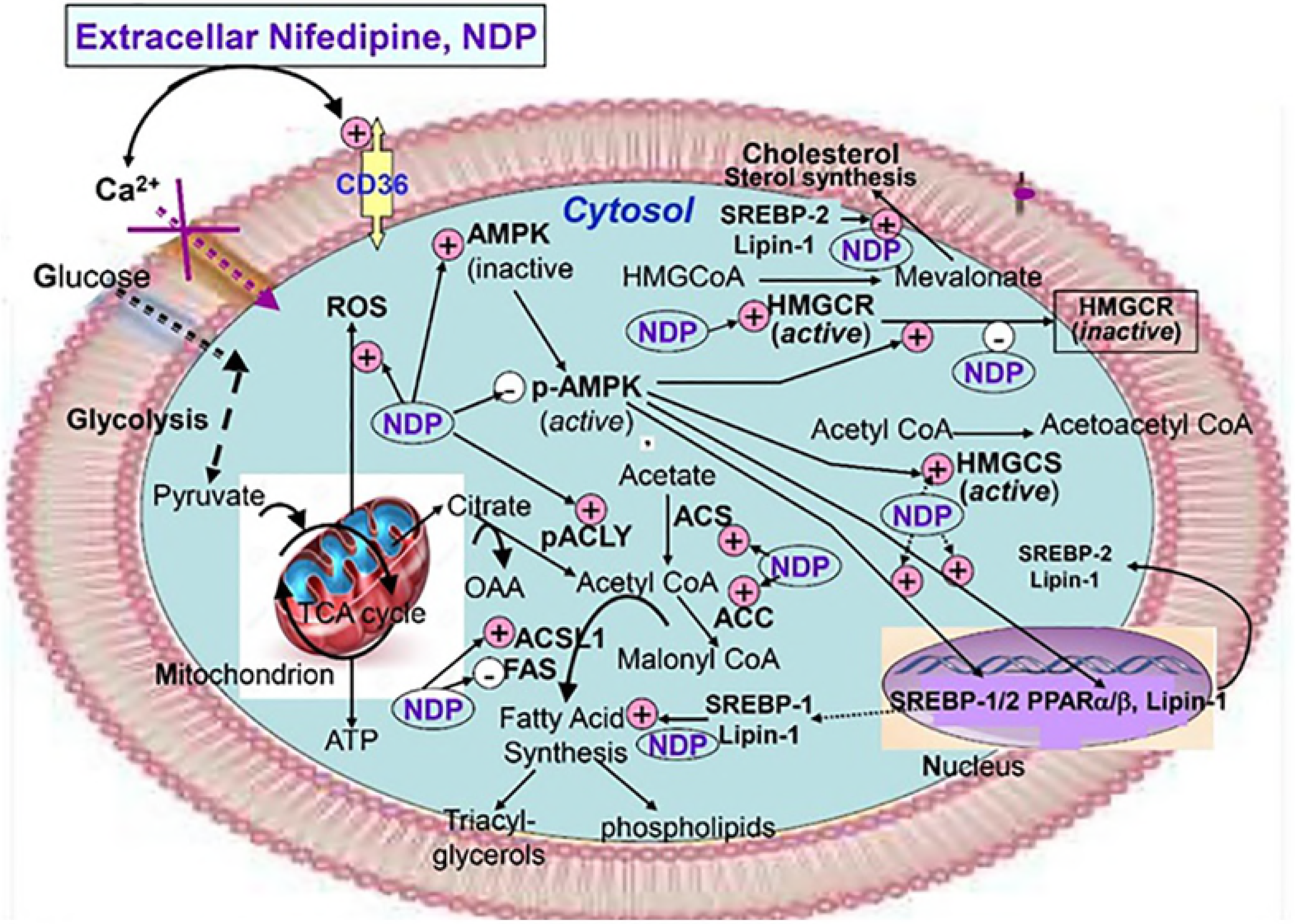

### Conclusions

Lipid accumulation in renal cells has been implicated in the pathogenesis of obesity-related kidney disease; however, nifedipine-induced renal lipotoxicity has never been cited. To our knowledge, this is the first report demonstrating nifedipine-induced lipid accumulation in the kidney. Renal lipotoxicity is a surrogate marker for renal failure or renal fibrosis. Prolonged and chronic treatment with nifedipine may induce or potentiate acute kidney injury (AKI), which has been etiologically considered to be closely related to CKD [46].

Although prior research demonstrated that nifedipine inhibited lipogenesis via activation of the LKB1-AMPK pathway, our findings indicate a lipogenic effect of nifedipine in renal cells. In brief, nifedipine induces the accumulation of cytosolic FFA, triggering the translocation of FFA into mitochondria to stimulate the production of ROS, leading to mitochondrial dysfunction (due to loss of cristae) and reducing β-oxidation with subsequent cellular lipid accumulation and kidney cell injury. Simultaneously, p-AMPK activity is inhibited (and, conversely, mTOR is upregulated), which may trigger the transcription of *SREBP-1/2* and *lipin-1*. Overall, in this work we have confirmed the mechanism of action involved in the renal lipotoxicity induced by nifedipine therapy, which can be elicited via the AMPK-SREBP-1/2 pathway. Our findings suggest possible novel strategies for prevention of such renal lipotoxicity through the administration of certain antioxidant therapies or agents targeting mitochondria.

## Acknowledgments

We wish to thank Professor Robert Y. Peng for editing and reviewing the whole manuscript. We are grateful to Ms. Chin-Yuan Chung for technical assistance.

